# MicroRNAs Regulate the Wnt/Ca^2+^ Signaling Pathway to Promote the Secretion of Insulin in Pancreatic Nestin-Positive Progenitor Cells

**DOI:** 10.1101/003913

**Authors:** Chunyu Bai, Xiangchen Li, Yuhua Gao, Taofeng Lu, Kunfu Wang, Qian Li, Hui Xiong, Jia Chen, Ping Zhang, Wenjie Wang, Tingting Sun, Zhiqiang Shan, Sun Bo, Pei Pei, Changqing Liu, Weijun Guan, Yuehui Ma

## Abstract

MicroRNAs (miRNAs) are small noncoding RNAs that bind to the 3′-UTR of mRNAs and function mainly in post-transcriptional regulation. MiRNAs have been implicated to play roles in organ development, including that of the pancreas. Many miRNAs, such as miR-375, miR-124, miR-7, miR-21 and miR-221, have been shown to regulate insulin production as well as insulin secretion. However, it is not known whether miRNAs can regulate insulin secretion via the control of intracellular Ca^2+^ in pancreatic beta cells. In this research, expression profiles of miRNAs and mRNAs were investigated using RNA-sequencing and microarray analysis in chicken pancreatic nestin-positive progenitor cells and differentiated pancreatic beta cells. A number of miRNAs were up-regulated after differentiation of progenitors into beta cells, which regulate cell signaling pathways to control cell function. miR-223 and miR146a were shown to promote insulin secretion from pancreatic beta cells by regulating the concentration of intracellular Ca^2+^ via the down-regulation of their target genes.

## Introduction

Pancreatic beta cells have well-tuned machinery for sensing glucose and secreting insulin. Type 2 glucose transporters mediate the uptake of glucose in beta cells, and subsequent glycolysis then leads to an increase in the ATP/ADP ratio. High ATP levels close potassium channels, preventing the exit of K^+^ ions. An excess positive charge of K^+^ ions in the cytosol depolarizes the membrane, which in turn results in the opening of voltage-gated calcium channels. Calcium ions then flow into the beta cell, inducing the exocytosis of insulin-containing granules (Goke 2008; Leibiger et al. 2008; Nolan and Prentki 2008).

The Wnt signaling pathway is involved in many physiological and pathophysiological activities. Wnt signaling is complex since the signals are transduced through several different pathways, depending on the Wnt type and the tissue analyzed (Pandur et al. 2002; Nelson and Nusse 2004; Dejmek et al. 2006). Most Wnts can be separated into two classes: canonical Wnts, which associate with LRP5/6/frizzled (Fzd) receptor complexes to stabilize b-catenin, which then activates Tcf/LEF-family transcription factors, and non-canonical Wnts, which associate with Fzd and other receptors, such as Ror2, to activate small GTPases and protein kinases to regulate cell migration and tissue polarity (Elizalde et al. 2011). Several non-canonical Wnt pathways have been reported, including the Wnt/Ca^2+^ signaling, Wnt planer cell polarity, Wnt-JNK signaling, Wnt/Ror receptor, Wnt-GSK3MT and Wnt-aPKC pathways (Semenov et al. 2007; Qiu et al. 2011; Thrasivoulou et al. 2013). Many in vitro and in vivo studies have shown that several components of the WNT pathway are involved in pancreatic beta cell proliferation (Rulifson et al. 2007; Shu et al. 2008), normal cholesterol metabolism, glucose-induced insulin secretion and the production of the incretin hormone, glucagon-like peptide-1 (Fujino et al. 2003; Yi et al. 2005). The Wnt/Ca^2+^ signaling pathway plays a crucial role in the development of the embryo and various organs. Signaling by Wnt-5a can trigger the non-canonical Wnt/Ca^2+^ pathway, leading to activation of the intracellular Wnt/Ca^2+^ signaling pathway (Dejmek et al. 2006; Witze et al. 2013). However, the function of the Wnt/Ca^2+^ signaling pathway in insulin secretion from pancreatic beta cells remains unknown.

A major group of endogenous small noncoding ribonucleotides (18-24 nt) called microRNAs (miRNAs), present in animals and plants, has been shown to play important roles in the regulation of gene expression at the post-transcriptional level (Bartel 2004; Tang et al. 2007; Tang et al. 2009). Specific miRNAs act in endocrine tissues, for example, miR-375 is highly expressed in the endocrine pancreas (Poy et al. 2004; Avnit-Sagi et al. 2009), and loss of its function disrupts islet morphogenesis and endocrine cell differentiation (Poy et al. 2009). However, miR-375 does not influence intracellular Ca^2+^ signaling to regulate insulin secretion from pancreatic beta cells. Here, we characterized miR-1552 and miR-489 expression and identified their functions in pancreatic beta cells. We show that miR-1552 and miR-489 directly block Wnt/Ca^2+^ signaling by repressing Wnt5a mRNA expression and by influencing the concentration of intracellular Ca^2+^, thereby regulating insulin secretion.

## Results

### Characteristics of Chicken Pancreatic Nestin-Positive Progenitor Cells (CPNPCs)

In pancreatic islets, only a small subset of cells express nestin, and these cells are proposed to represent precursors of differentiated pancreatic endocrine cells (Hunziker and Stein 2000). These nestin-positive cells are called pancreatic stellate cells and are proposed to play an important role in the growth and maintenance of islets (Lardon et al. 2002; Selander and Edlund 2002; Treutelaar et al. 2003). Chicken pancreatic nestin-positive progenitor cells (CPNPCs) were isolated from embryonic day 16 chicks according to the method of Zulewski (2001). In primary culture, CPNPCs formed colonies 4 days after seeding. Many erythrocytes were mixed with the CPNPCs; however, the CPNPC population was purified from hemocytes and other cells using flow cytometry (Fig. 1 A). The cells showed cluster formation and maintained their growth capacity. The colony-forming units were 42.65±2.54%, 40.6±1.35% and 43.57±3.34% for passages 10, 20 and 30, respectively using colony-forming cell assay, demonstrating the capacity of the cultured CPNPCs to self-renew. Pancreatic and duodenal homeobox 1 gene (Pdx1) is also known as insulin promoter factor 1 (Ipf1) or islet/duodenum homeobox 1 (IDX1). In the adult mouse, it is selectively expressed in islet beta cells, where it binds to and regulates the insulin promoter (Ohlsson et al. 1993). Pdx1, a transcription factor, is recognized as a marker for pancreatic precursors in pancreatic development (Pennarossa et al. 2013). We examined cells for nestin and Pdx-1 expression using immunofluorescence and flow cytometry, and the results demonstrated that cells in cluster formations were double-positive for nestin and Pdx1 (Fig. 1 B and Fig. 1 C).

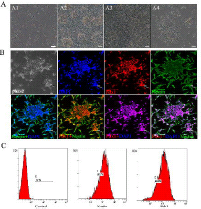

### Differentiation of CPNPCs towards islet beta cells

Nicotinamide, a poly (ADP-ribose) synthetase inhibitor, has been reported to prevent the development of diabetes in animal models, and to promote the proliferation and differentiation of islet beta cells (Uchigata et al. 1983; Otonkoski et al. 1993). We used 10 mM nicotinamide to induce the differentiation of islet beta cells from CPNPCs, and we then observed changes in the transcription of the islet hormone genes, Pdx1, ISL1, INS and MYT1 (Fig. 2 C and F) Insulin release is an important characteristic of islet beta cells. Glucose-induced insulin secretion demonstrated that the CPNPC-derived clusters could release insulin at glucose concentrations of 5.5 mM and 17.5 mM. Non-differentiated CPNPCs did not release insulin.(Fig. 2 A and B) Type 2 glucose transporters mediate the uptake of glucose in beta cells and subsequent glycolysis leads to an increase in the ATP/ADP ratio. High ATP levels close potassium channels, preventing the exit of K^+^ ions. The excess of positive charge of the K^+^ ions in the cytosol depolarizes the membrane, which in turn results in the opening of voltage-gated calcium channels. Calcium ions (Ca^2+^) then flow inside the cell, inducing the exocytosis of insulin-containing granules. The concentration of intracellular calcium ions was analyzed using flow cytometry and video observation after Fluo-4 AM labeling. The fluorescence value of Ca^2+^ was significantly up-regulated after 17.5mM glucose-induction using observation of confocal optical system and flow cytometry. (Supplementary video and Fig.2 D)

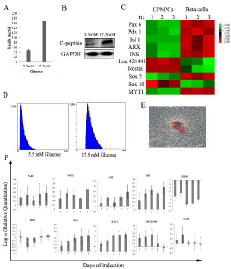

### Differentially Expressed Genes in CPNPCs and islet beta cells

The relative expression levels of mRNAs in CPNPCs and islet beta cells were evaluated using microarray analysis. Significantly differentially expressed genes were selected and analyzed for their roles in biological processes, molecular function or as cellular components. Some signaling pathways were discovered via Gene Ontology (GO) analyses, including pathways for notch, PDGF, TFGF-beta and Wnt. The Notch signaling pathway plays critical roles in many developmental processes, influencing differentiation, proliferation and apoptosis (Greenwald 1998; Artavanis-Tsakonas et al. 1999). Endocrine differentiation in the early embryonic pancreas is regulated by Notch signaling. Activated Notch signaling maintains pancreatic progenitor cells in an undifferentiated state, whereas suppression of Notch leads to endocrine cell differentiation (Xu et al. 2006). The myelin transcription factor 1 (MYT1), a zinc finger DNA-binding protein and a target of Notch signaling is expressed at low levels in CPNPCs. However, the Notch signaling pathway was inhibited as CPNPCs differentiated towards islet beta cells, and expression of the MYT1 gene was increased in islet beta cells (Fig. S1 and Fig. S2). The PDGF signaling pathway promotes proliferation, survival and migration in diverse cell types (Schlessinger 2000). Previous studies have demonstrated that activation of PDGF signaling stimulates DNA synthesis in cultured islets (Welsh et al. 1990; Su et al. 2005) and regulates age-dependent proliferation of islet beta cells (Chen et al. 2011). The expression of platelet-derived growth factor subunit A (PDGFA) was up-regulated after differentiation of islet beta cells, indicating that the PDGF signaling pathway may regulate the proliferation of islet beta cells after differentiation (Fig. S3 and Fig. S4). Wnt family members are secreted proteins that signal through the frizzled superfamily of G protein-coupled receptors, and are involved in many physiological and pathophysiological activities. The Wnts were divided into two classes according to their specific receptors: the “canonical” or the “non-canonical” signaling pathways (McDonald and Silver 2009; Gregory et al. 2010). Non-canonical Wnt pathways include the planar cell polarity pathway and the Ca^2+^ pathway. The non-canonical Wnt signaling pathway plays an important role in insulin secretion from islet beta cells. The expression level of Wnt5A was down-regulated in differentiated islet beta cells (Fig. 3). This could inhibit the Ca^2+^ pathway to maintain the concentration of intracellular Ca^2+^ and promote the release of insulin.

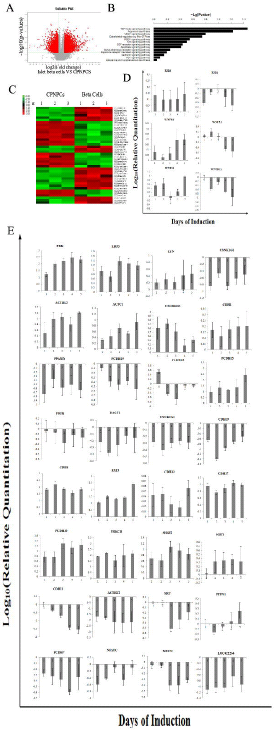

### Differential expression profile of microRNAs in CPNPCs and islet beta cells

Many miRNAs can regulate insulin secretion from beta cells, such as miR-375, miR-124a and let-7 (Krek et al. 2005; Lovis et al. 2008). MiRNAs in animals exhibit tissue-specific or developmental-stage-specific expression, indicating that they play important roles in many biological processes (Lagos-Quintana et al. 2001; Lau et al. 2001). However, there has been no report describing the functions of miRNAs in the differentiation of islet beta cells from pancreatic progenitor/stem cells. Therefore, we applied a deep sequencing approach to reveal the differential expression profile of miRNAs in CPNPCs and islet beta cells. The differential expression profile of miRNAs is shown in Table 1 and Fig. 4A. Nine miRNAs (indicated in red in Table 1) were up-regulated after differentiation of islet beta cells. The target genes of these differentially expressed miRNAs were predicted using Targetscan tools (http://www.targetscan.org/) and analyzed using Venny tools (http://bioinfogp.cnb.csic.es/tools/venny/index.html). The target genes of differentially expressed miRNAs and differentially expressed genes between CPNPCs and pancreatic beta cells were assessed in a Venn diagram. This analysis identified 727 genes in the intersection of the Venn diagram, and these genes were then investigated by GO and Pathway analysis (Fig. 4 B, C and D). Pathway analysis demonstrated that miRNAs could regulate the signaling pathways that control the differentiation of islet beta cells from CPNPCs, such as Notch, PDGF, Wnt, and Cadherin signaling pathways. MiRNAs that were up-regulated during the differentiation of islet beta cells could down-regulate crucial components of these signaling pathways to promote the differentiation of islet beta cells (Fig. 4 E).

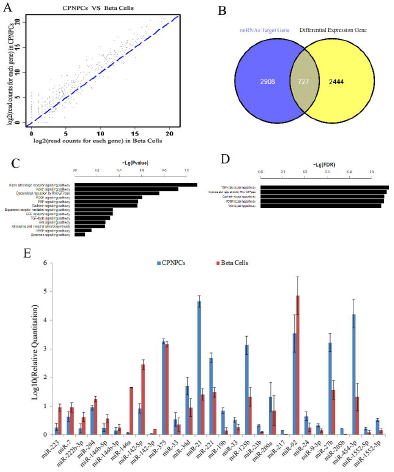

### MicroRNAs regulated the concentration of intracellular Ca^2+^ to promote insulin secretion

Previous studies demonstrated that miRNAs can regulate the differentiation of islet beta cells and the secretion of insulin from the pancreas (Poy et al. 2004; Kredo-Russo et al. 2012; Klein et al. 2013). miR-375 was identified to regulate the secretion of insulin. One of the targets of miR-375 was identified as myotrophin, a protein implicated in exocytosis, but not intracellular Ca^2+^ signaling. In our study, miRNAs were observed from their differential expression profiles to regulate the Wnt5a-controlled secretion of insulin via intracellular Ca^2+^. The miRNAs, gga-miR-223 and gga-miR-146a, were overexpressed using microRNA mimics to inhibit the target gene expression (Wnt5a) in beta cells, in opposite, the Wnt5a were overexpressed in beta cells (Fig. 5A and B). The concentration of intracellular Ca^2+^ and the secretion of insulin were analyzed using the non-invasive micro-test technique (Kuhtreiber and Jaffe 1990) and western blotting. The Wnt5a can control directional movement to mobilize the Ca^2+^ signal in the cell, activating actomyosin contraction and adhesion disassembly for membrane retraction (Witze et al. 2013). The results demonstrated that overexpression of miR-223 and miR-146a could increase the concentration of intracellular Ca^2+^ to promote the insulin secretion in beta cells, the counter-productive results were overexpression of Wnt5a (Fig. 5 D, E and F).

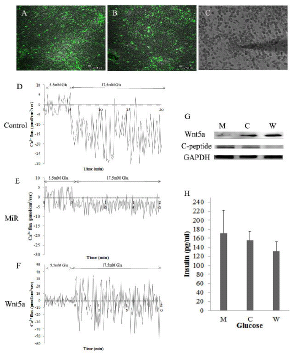

## Discussion

Many miRNAs have previously been reported to regulate insulin secretion and development of the pancreas, such as miR-375 (Poy et al. 2004; Poy et al. 2009), miR-30d (Tang et al. 2009), miR-124a (Baroukh et al. 2007), miR-7 (Kredo-Russo et al. 2012), miR-21 (du Rieu et al. 2010), miR-221 (Farrell et al. 2013) and miR-19b (Zhang et al. 2011). However, there has been no demonstration of miRNAs that can promote insulin secretion through the regulation of intracellular Ca^2+^ concentration. Here, we show that gga-miR-223 and gga-miR-146a can regulate the concentration of intracellular Ca^2+^ to increase insulin secretion through suppression of the Wnt/Ca^2+^ signaling pathway. The Wnt/Ca^2+^ pathway is stimulated by certain Wnt ligands that activate Frizzled (Fz) receptors and involves the calcium-responsive enzymes, PKC and calcium/calmodulin-dependent protein kinase II (CamKII) (Kuhl et al. 2000a; Kuhl et al. 2000b). Wnt5a has been proposed to exert its effects via the Wnt/Ca^2+^ pathway, based on results obtained in both Xenopus and mammalian cells (Saneyoshi et al. 2002). Wnt5a can release intracellular Ca^2+^ to promote edge polarization of the cell surface (Witze et al. 2013).

Mitochondria and Ca^2+^ influx regulate glucose stimulated insulin secretion from pancreatic beta cells. Glucose is transported into the cytosol and metabolized into pyruvate. Mitochondria then metabolize pyruvate to produce adenosine triphosphate (ATP) and other metabolic products via the tricarboxylic acid cycle (Maechler and Wollheim 2001; Pertusa et al. 2002). ATP can trigger closure of ATP-sensitive potassium channels (KATP channels) contributing to membrane depolarization. Besides acting on the KATP channels, ATP interacts with other products of mitochondrial metabolism that increase the production of IP3 (inositol 1,4,5-trisphosphate) (Liu et al. 1996), which in turn mobilizes Ca^2+^ from the endoplasmic reticulum (Dyachok and Gylfe 2004). Ca^2+^ mobilization activates Ca^2+^ channels on intracellular stores (Liu and Gylfe 1997), and this current cooperates with the closure of KATP channels, resulting in membrane depolarization and activation of voltage-dependent Ca^2+^ channels. Ca^2+^ channels are considered the most important means of Ca^2+^ entry and this Ca^2+^ signal controls insulin secretion from pancreatic beta cells (Valdeolmillos et al. 1992). Wnt5a and the Wnt/Ca^2+^ signaling pathway also activates edge polarization of the cell surface, which reduces the concentration of intracellular Ca^2+^ to restrict insulin secretion from pancreatic beta cells.

The high level of expression of miRNAs, including gga-miR-7, gga-miR-223, gga-miR-222b, gga-miR-204, gga-miR-146ab and gga-miR-142, in pancreatic beta cells differentiated from CPNPCs, suggests that they play important roles in this differentiation. Bioinformatic predictions and GO analysis suggest that their target genes play a role in pancreas biology. We focused on the regulation of Wnt5a by miRNAs because of our bioinformatics analysis and because Wnt5a is a trigger for Wnt/Ca^2+^ signaling to regulate the concentration of intracellular Ca^2+^. Over-expression of miRNA-mimics and Wnt5a demonstrated that Wnt5a was an agonist for decreasing intracellular Ca^2+^ concentration. Over-expression of Wnt5a can disturb pancreatic beta cell function through intracellular Ca^2+^.

### Materials and Methods

Research into human pancreas development has been restricted by ethical constraints and access to tissue, particularly during the first trimester, placing understandable reliance on data from other species, such as mouse, rat, frog and chick. The chicken is a classic model of vertebrate developmental biology and medicine that has been used for many decades (Brown et al. 2003). So, the chicken was used as an animal model in our research. Chicken embryos were provided by the chicken breeding farm of the Chinese Academy of Agricultural Sciences, Beijing, China. Animal experiments were performed in accordance with the guidelines established by the Institutional Animal Care and Use Committee of the Chinese Academy of Agricultural sciences.

### Isolation and purification of nestin-positive stem cells from chick embryos

Nestin-positive stem cells were isolated from 15-day-old chick embryonic pancreas by the collagenase digestion method of Joel F. Habener (Zulewski et al. 2001) and cultured in advanced RPMI 1640 media (11.1 mmol/L glucose) supplemented with 10% fetal bovine serum, 1 mmol/L sodium pyruvate, 20 ng/mL bFGF, 20 ng/mL EGF and 71.5 mmol/L β-mercaptoethanol. The fraction containing the majority of nestin-positive stem cells was obtained after purification through flow cytometry (FCM). Immunofluorescence was used to analyze nestin and Pdx1 in stem cells.

### Differentiation of nestin-positive stem cells towards pancreatic beta cells

For differentiation towards endocrine pancreatic phenotypes, clonal cell masses were picked and plated on 24-well cell culture plates, then cultured for 7 days in Dulbecco’s modified Eagle’s medium/F12 containing glucose (11.1 mM) and a cocktail of several growth factors, including 10 mM nicotinamide, 100 pmol/L HGF and 2 nmol/L active-A. To test whether insulin release from differentiated cells was glucose-dependent, two glucose concentrations (5.5 mM and 25.5 mM) were used (Otonkoski et al. 1988). Insulin release and c-peptide were measured using the Insulin ELISA Kit and western blotting, respectively. The calcium ion concentration of cells was analyzed using Flou-4/AM.

### Total RNA isolation, small RNA library construction, and Solexa sequencing

Total RNA samples were size fractionated by 15% polyacrylamide gel electrophoresis, and the 16-30 nt fraction was collected. 5′ and 3′ RNA adaptors were ligated to the RNA, and RNAs of 64-99 nt were isolated by gel elution and ethanol precipitation. Polymerase chain reaction (PCR) products were purified, and small RNA libraries were sequenced using an Illumina Genome Analyzer. Sequencing was carried out at the Shanghai Biotechnology Corporation (SBC, Shanghai, China).

### Microarray analysis

RNA samples from three independent experiments were hybridized to a chicken gene expression microarray (Agilent, 4*44K, Design ID 026441) following the manufacturer instructions. For each sample, the background was removed and raw data were normalized using the MAS 5.0 algorithm (Gene Spring Software 11.0; Agilent technologies). Heat maps were generated using Java Treeview software. Microarray data and RNA-seq data have been deposited in the NCBI’s Gene Expression Omnibus public database (accession number SRP034850).

### Gene Ontology (GO) category and pathway analysis

The PANTHER Classification System (http://www.pantherdb.org/) was used to interpret the biological effect of mRNAs and the target genes of miRNAs. GO categorization of biological processes for the differentially expressed and target genes was performed using the Gene Ontology project (background: ftp://ftp.pantherdb.org/sequence_classifications/current_release/PANTHER_Sequence_Classification_files/PTHR8.1_chicken). The Kyoto Encyclopedia of Genes and Genomes (KEGG) genome database was used to identify significant gene pathways during the differentiation of nestin-positive stem cells.

### Over-expression of miRNAs and Wnt5a in pancreatic beta cells

miRNA mimics and a control sequence were synthesized by GenePharma (Shanghai, China) and transfected into pancreatic beta cells using RNA-Mate reagent. Real-time PCR and western blot analyses were used to detect the expression of miRNAs and their target genes. The coding regions of Wnt5a were PCR-amplified and cloned into a lentiviral vector. This vector was then incorporated into a lentivirus and transfected into HEK-293T cells. The lentivirus also expressed GFP as a marker for monitoring infection efficiency.

